# Recombinant ACE2 Expression is Required for SARS-CoV-2 to Infect Primary Human Endothelial Cells and Induce Inflammatory and Procoagulative Responses

**DOI:** 10.1101/2020.11.10.377606

**Authors:** Jonas Nascimento Conde, William Schutt, Elena E. Gorbunova, Erich R. Mackow

## Abstract

SARS-CoV-2 causes COVID-19, an acute respiratory distress syndrome (ARDS) characterized by pulmonary edema, viral pneumonia, multiorgan dysfunction, coagulopathy and inflammation. SARS-CoV-2 uses angiotensin-converting enzyme 2 (ACE2) receptors to infect and damage ciliated epithelial cells in the upper respiratory tract. In alveoli, gas exchange occurs across an epithelial-endothelial barrier that ties respiration to endothelial cell (EC) regulation of edema, coagulation and inflammation. How SARS-CoV-2 dysregulates vascular functions to cause ARDS in COVID-19 patients remains an enigma focused on dysregulated EC responses. Whether SARS-CoV-2 directly or indirectly affects functions of the endothelium remains to be resolved and critical to understanding SARS-CoV-2 pathogenesis and therapeutic targets. We demonstrate that primary human ECs lack ACE2 receptors at protein and RNA levels, and that SARS-CoV-2 is incapable of directly infecting ECs derived from pulmonary, cardiac, brain, umbilical vein or kidney tissues. In contrast, pulmonary ECs transduced with recombinant ACE2 receptors are infected by SARS-CoV-2 and result in high viral titers (∼1×10^7^/ml), multinucleate syncytia and EC lysis. SARS-CoV-2 infection of ACE2-expressing ECs elicits procoagulative and inflammatory responses observed in COVID-19 patients. The inability of SARS-CoV-2 to directly infect and lyse ECs without ACE2 expression explains the lack of vascular hemorrhage in COVID-19 patients and indicates that the endothelium is not a primary target of SARS-CoV-2 infection. These findings are consistent with SARS-CoV-2 indirectly activating EC programs that regulate thrombosis and endotheliitis in COVID-19 patients, and focus strategies on therapeutically targeting epithelial and inflammatory responses that activate the endothelium or initiate limited ACE2 independent EC infection.

**Importance:** SARS-CoV-2 infects pulmonary epithelial cells through ACE2 receptors and causes ARDS. COVID-19 causes progressive respiratory failure resulting from diffuse alveolar damage and systemic coagulopathy, thrombosis and capillary inflammation that tie alveolar responses to EC dysfunction. This has prompted theories that SARS-CoV-2 directly infects ECs through ACE2 receptors, yet SARS-CoV-2 antigen has not been co-localized with ECs and prior studies indicate that ACE2 co-localizes with alveolar epithelial cells and vascular smooth muscle cells, not ECs. Here we demonstrate that primary human ECs derived from lung, kidney, heart, brain and umbilical veins require expression of recombinant ACE2 receptors in order to be infected by SARS-CoV-2. However, SARS-CoV-2 lytically infects ACE2-ECs and elicits procoagulative and inflammatory responses observed in COVID-19 patients. These findings suggest a novel mechanism of COVID-19 pathogenesis resulting from indirect EC activation, or infection of a small subset of ECs by an ACE2 independent mechanism, that transform rationales and targets for therapeutic intervention.

---

SARS-CoV-2 predominantly infects the epithelium of upper and lower airways causing pulmonary pathology and ARDS^(1)^. COVID-19 is characterized by progressive respiratory failure resulting from diffuse alveolar damage, inflammatory infiltrates, endotheliitis and pulmonary and systemic coagulopathy forming obstructive microthrombi with multiorgan dysfunction^(1–3)^. Collectively these findings indicate that initial pulmonary epithelial infection leads to COVID-19 vasculopathy with featured alveolar endothelial cell (EC) dysfunction playing a key role in anomalous vascular leakage, coagulation and inflammation. In COVID-19 patients procoagulative responses are associated with altered von Willebrand factor (vWF) and thrombomodulin expression, and the induction of proinflammatory cytokines (IL1, IL6, TNFα) that further implicate activation of the endothelium in myocarditis and vasculopathy^(1–4)^.

Despite coagulopathy and capillary inflammation in COVID-19 patients, it is unclear whether ECs are directly infected by SARS-CoV-2 or whether EC activation is an indirect response to primary alveolar epithelial cell damage and inflammatory responses^(1–3)^. SARS-CoV-2 infects cells by attaching to human ACE2 receptors^(5–7)^. Rationales for SARS-CoV-2 directly infecting ECs originated from prothrombotic findings, endotheliitis, protective ACE2 functions and reports that ECs express cellular ACE2 receptors^(8–10)^. However, several studies demonstrate that in the vasculature ACE2 is confined to the tunica media, co-localizing with smooth muscle actin, not the endothelium^(11–14)^. CDC analysis of COVID-19 patient tissues indicates that SARS-CoV-2 is detectable in airways, pneumocytes, alveolar macrophages and lymph nodes, but not in ECs or other extrapulmonary tissues^(1)^. In retrospect, there is minimal data supporting SARS-CoV-2 infection of ECs and no immunohistochemical studies demonstrating the co-localization of SARS-CoV-2 antigens with EC markers in pulmonary or renal tissues, which express ACE2 on adjacent epithelial cells. Nearly all studies reference electron microscopy data displaying two potential SARS-CoV-2 particles^(3, 15)^, that instead of virus have been implicated as being ER vesicles^(16)^.

Nonetheless, pathologic findings in COVID-19 patients demonstrate the dysregulation of EC functions^(17)^, however, the mechanism(s) of endothelial damage and activation in SARS-CoV-2 directed coagulopathy and inflammation remain to be revealed^(2, 4)^. Our initial studies were predicated on ACE2 receptors directing SARS-CoV-2 infection and dysregulation of normal EC functions. We critically analyzed SARS-CoV-2 infection of primary human ECs derived from lung, heart, kidney, brain and umbilical veins (S1-methods). Remarkably we found that SARS-CoV-2 failed to infect primary human ECs derived from any organ. In contrast to the complete infection of VeroE6 cells, no SARS-CoV-2 infected ECs were detected, by N or Spike antigen immunostaining, at any multiplicity of infection or plating cell density (Figure 1A). Consistent with this both ACE2 RNA and protein, found in VeroE6 and Calu3 cells, were undetectable in ECs (Figure 1B,C), and no viral progeny was detected in the supernatants of SARS-CoV-2 infected human ECs (1-3 dpi)(Figure 1G).

**Figure 1.**
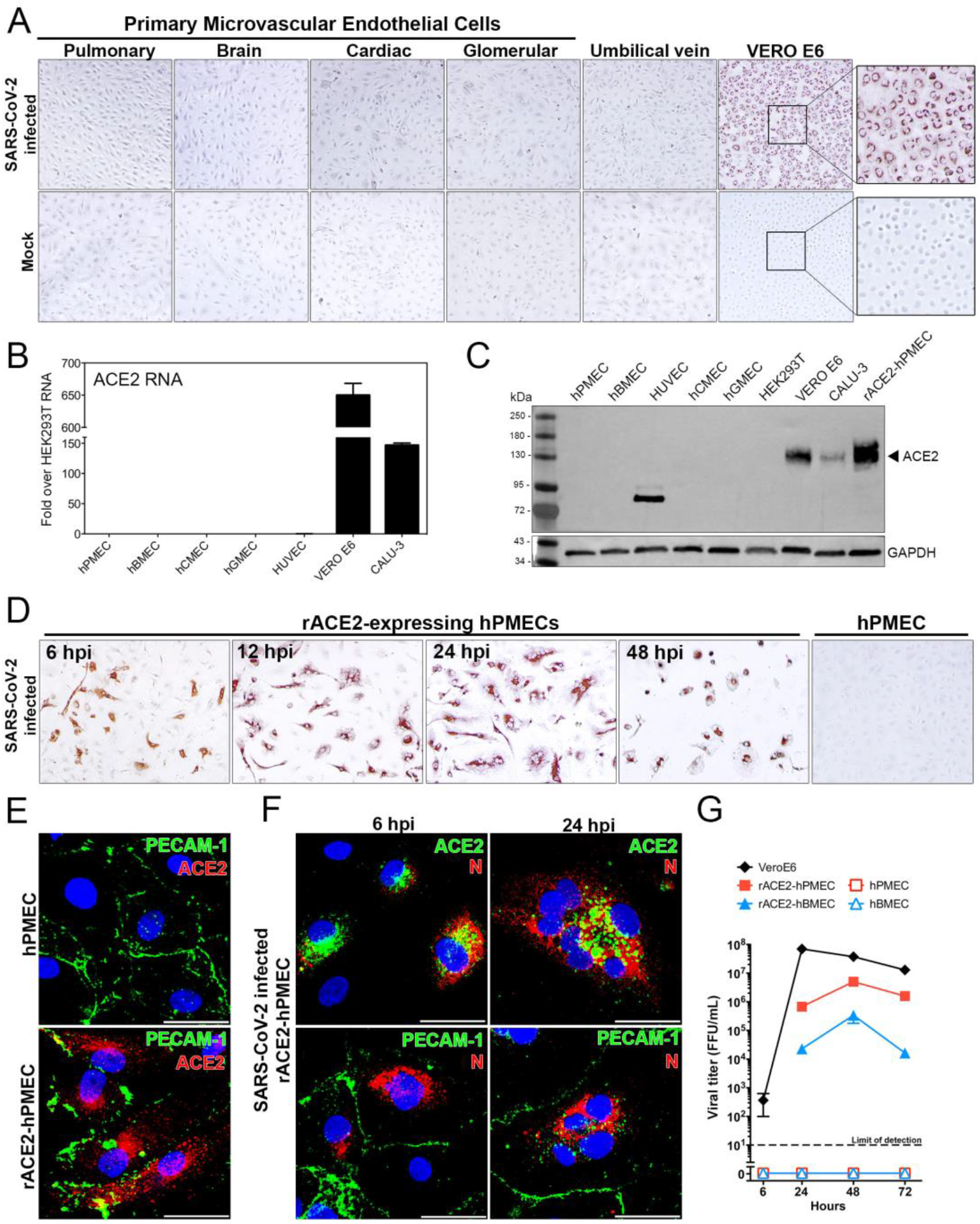
SARS-CoV-2 Fails to Infect Primary Human Endothelial Cells without rACE2 Expression. Primary human microvascular endothelial cells from pulmonary (hPMECs), brain (hBMECs), cardiac (HCMECs), glomerular (hGMECs) or umbilical vein (HUVECs), or VeroE6 cells were mock or SARS-CoV-2(strain WA) infected (MOI 10) and 24 hpi **(A)** immunoperoxidase stained for nucleocapsid protein. Primary human ECs, VeroE6, HEK293T and Calu3 cells were analyzed by **(B)** qRT-PCR for ACE2 mRNA, and **(C)** by Western Blot for expressed ACE2. HUVECs contain a potential ACE2 truncation lacking the N-terminal SARS-CoV-2 binding domain. **(D)** Primary human pulmonary ECs lentivirus transduced to express recombinant ACE2 were infected with SARS-CoV-2 (MOI 1) for 6-48 hpi. **(E)** ACE2-hPMECs or WT-hPMECs were immunoperoxidase stained for nucleocapsid protein. hPMECs or rACE2-hPMECs were analyzed by IFA for the EC marker PECAM-1 and ACE2. **(F)** Following SARS-CoV-2 infection, rACE2-hPMECs were analyzed by IFA for coexpressed ACE2 and nucleocapsid protein (N) or PECAM-1 expression. Bars represent 50 µm. **(G)** Supernatants of Sars-CoV-2 infected (MOI 1) WT hPMECs, hBMECs, rACE2-hPMECs, rACE2-hBMECs and VeroE6 cells were titered 2-72 hpi (Limit of detection <10 FFUs/ml).

To determine whether SARS-CoV-2 infection of ECs is receptor restricted, we lentivirus transduced primary human pulmonary and brain ECs to express ACE2 and evaluated viral replication and protein expression. We found that expressing ACE2 in primary human ECs permitted SARS-CoV-2 to ubiquitously and productively infect ECs reaching viral titers of 1-3 x 10^7^ (1-3 dpi) (Figure 1D,G)(S1-methods). SARS-CoV-2 infection co-localized with ACE2 expressing ECs (Fig. 1E,F) and resulted in detectable N protein 4-6 hpi, and multinucleate syncytia and EC lysis 12-24 hpi (Figure 1D,F). Collectively, these findings demonstrate that primary human ECs lack ACE2 receptors required for SARS-CoV-2 infection, but express proteases essential for SARS-CoV-2 infection. These findings suggest that SARS-CoV-2 may cause procoagulative endotheliitis through indirect EC dysregulation mechanisms or as a result of ACE2 independent, or induction directed, infection of a small number of activated ECs.

The potential for damage, inflammation or activation to conditionally permit SARS-CoV-2 infection of a small EC subset^(12, 18, 19)^, prompted us to analyze cellular responses that may contribute to COVID-19 pathogenesis. We analyzed transcriptional responses of ACE2-expressing ECs to SARS-CoV-2 infection and found significant changes in mRNAs that regulate coagulation and inflammation from 6-24 hrs (S1-methods) including: 2-3-fold decreases in PAI-1, anti-thrombin and Factor VIII; and increases in tissue factor (24-fold), thrombomudulin (TM), 6-fold), vWF (3-fold), thrombin receptors (PAR1/3 3-fold), EGR-1 (37-fold), E-selectin (600-fold), IL-1β (28-fold), IL-6 (12-fold) and TNFα (160-fold)^(20, 21)^ (Figure 2A). SARS-CoV-2 selectively induced thrombomodulin in infected rACE2-hPMECs, with TM internally co-localized with viral N protein (Figure 2B), suggesting the potential for SARS-CoV-2 to sequester a coagulation inhibiting EC surface receptor^(20)^. However, a comprehensive assessment of coagulation and inflammatory mediators in SARS-CoV-2 infected epithelial and endothelial cells is required to fully understand EC activation events and complex coagulation factor and inflammatory responses that can be therapeutically targeted.

**Figure 2.**
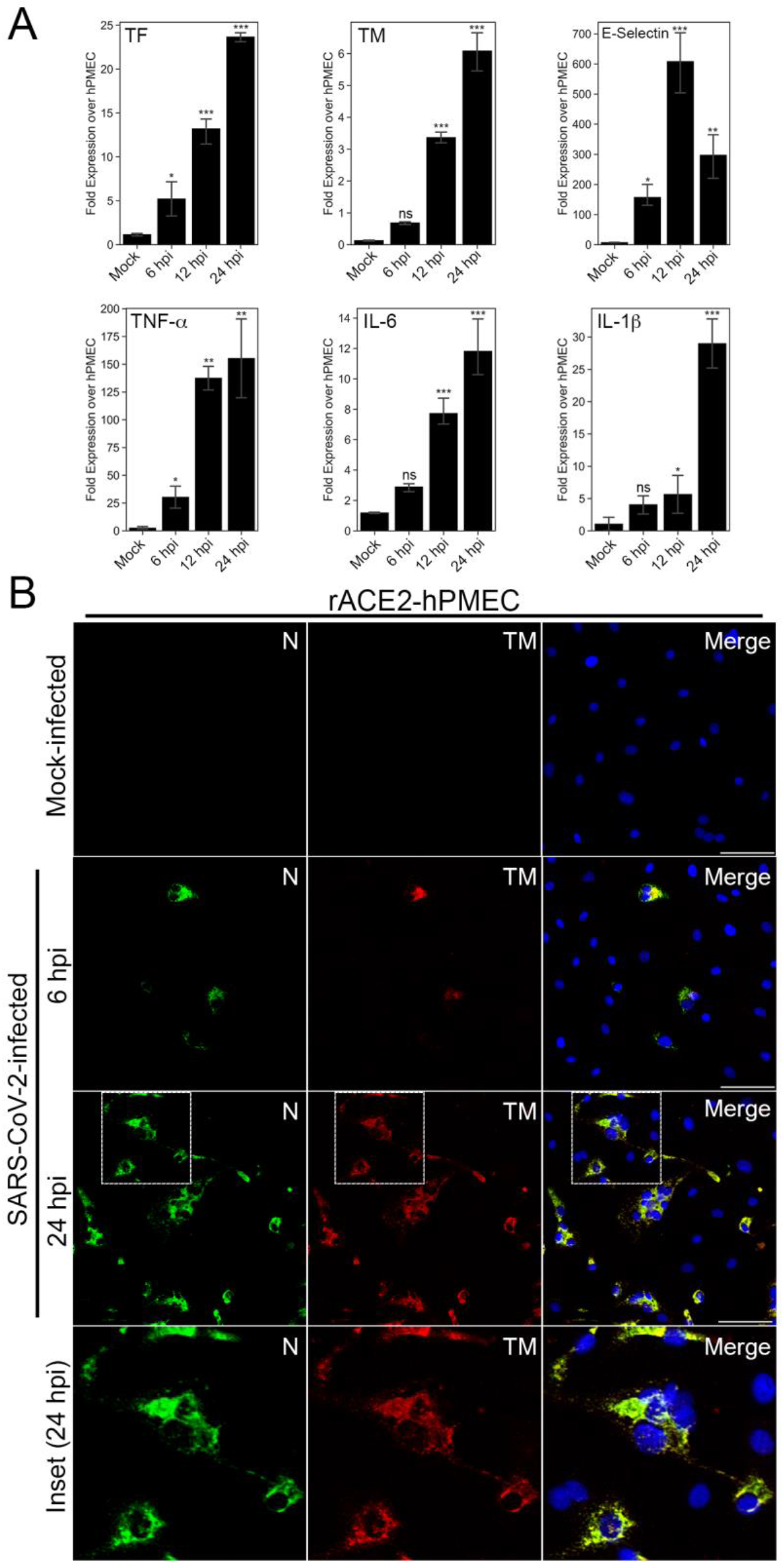
Recombinant ACE2 Expressing ECs Elicit Procoagulation and Inflammatory Responses. hPMECs expressing recombinant ACE2 (hPMEC-rACE2) were synchronously infected with SARS-CoV-2 and analyzed by qRT-PCR for changes in the mRNA levels of coagulation and inflammatory responses 6-24 hpi. **(A)** Levels of tissue factor (TF), thrombomodulin (TM), Tumor necrosis factor α (TNF α) Interleukin 6 (IL-6), IL-1β and E-selectin were found to increase dramatically in SARS-CoV-2 infected ECs. **(B)** The induction of TM in SARS-CoV-2 infected rACE2-hBMECs (MOI 1) was monitored by IFA of viral nucleocapsid (N) and cellular expressed thrombomodulin (TM) from 6-24 hpi.

Our findings indicate that the absence of ACE2 prevents SARS-CoV-2 infection of human ECs and suggests that ECs are not primary targets of SARS-CoV-2 infection in COVID-19 patients. Consistent with this, COVID-19 does not result in Ebola-like hemorrhagic disease that would likely result from lytic SARS-CoV-2 infection of ACE2-expressing ECs. The inability of SARS-CoV-2 to infect human ECs is supported by low ACE2 expression in the highly vascularized lower respiratory tract^(24)^, CDC and primary human EC infection findings^(1, 14, 24)^ and the presence of ACE2 in vascular smooth muscle and heart muscle cells^(11, 18, 23, 25)^, but not the EC lining of vessels^(12-14, 23)^. These findings support a secondary role of the endothelium, perhaps in response to epithelial cell damage and crosstalk, alveolar tissue factor/basement membrane exposure or inflammatory EC activation that directs a coagulative, endotheliitic state^(1, 3, 17, 22)^.

Our findings do not address whether SARS-CoV-2 infection of pulmonary epithelial cells permits SARS-CoV-2 to selectively infect or activate ECs. In the course of these experiments we tested, but were unable to define, conditions that permitted SARS-CoV-2 infection of pulmonary ECs (addition of Angiotensin II, activating AMP kinase, hypoxia, TNFα, IL-1β, IL6, bradykinin, endothelin-1). However, it remains conceivable that COVID-19 epithelial cell or immune cell responses selectively activate the endothelium^(2)^ and permit a subset of ECs to be infected over time^(19)^. Reported EC heterogeneity in response to acute lung injury^(19)^ supports the potential for infection of a subset of ECs, and one report suggests that 1/250 ECs are ACE2 positive, and that both SARS-CoV-2 and influenza virus increase the number of ACE2 positive ECs^(3)^. Yet SARS-CoV-2 infection of ACE2 expressing ECs remains to be demonstrated in COVID-19 patients and is at odds with current findings and additional studies indicating that ECs lack ACE2^(12–14, 23)^.

Consistent with COVID-19 disease we found that SARS-CoV-2 infection of ECs induces procoagulative and inflammatory mediators^(1–3, 17, 21)^. Our finding that the coagulation initiator, tissue factor, is highly induced, whereas the coagulation inhibitor thrombomodulin is induced and may be sequestered within ECs, provide potential thrombotic mechanisms, while findings of induced cytokines and E-selectin are consistent with inflammation and endotheliitis^(3, 20, 22, 26)^. These results rationalize a detailed analysis of EC expressed procoagulative and inflammatory factors and the potential role of targeting thrombomodulin, TNFα and E-selectin in resolving EC directed COVID-19 coagulation and inflammation^(3, 4, 20, 26)^. However, in the absence of EC infection, damage to alveolar epithelial cells may alone initiate coagulopathy through tissue factor, intra-alveolar fibrin deposition and common EC basement membrane exposure that triggers activation of the endothelium^(22, 27)^. In COVID-19 patients, EC damage and activation responses are also likely to be exacerbated by impaired ACE2 activity that increases the severity of ARDS, AngII directed EC damage, bradykinin directed permeability and inflammation, and the loss of protective anti-inflammatory Ang1-7 responses^(9, 25, 28–30)^. Overall, our data suggests that SARS-CoV-2 is likely to indirectly dysregulate EC functions, and this explains the absence of an acute lytic infection of ECs, and the chronic vascular disease process that over time evolves into an aberrant prothrombotic endotheliitis in COVID-19 patients. These findings focus strategies on therapeutically targeting epithelial and inflammatory responses that activate the endothelium or initiate limited ACE2 independent EC infection.

## Supplemental Material

The methods used within the studies describing cells, virus, SARS-CoV-2 infection, ACE2 lentivirus transduction, qRT-PCR analysis, Western Blotting, confocal Immunofluorescense and statistical analysis are presented in the supplemental section S1.

## Acknowledgements

This work was supported by a SARS-CoV-2 Seed Grant from Stony Brook University and funding from the Mathers Foundation and the National Institutes of Health NIAID R01AI12901004, R21AI13173902, R21AI15237201.We thank Ken Kaushansky, Berhane Ghebrehiwit and the SARS-CoV-2 SBU research group of Patrick Hearing, Janet Hearing, Nancy Reich and Hwan Kim for helpful input, discussions and critical review of the manuscript.

